# Neural correlates of interpersonal space permeability and flexibility in autism spectrum disorder

**DOI:** 10.1101/2020.10.14.339291

**Authors:** Claudia Massaccesi, Alexander Groessing, Lisa A. Rosenberger, Helena Hartmann, Michela Candini, Giuseppe di Pellegrino, Francesca Frassinetti, Giorgia Silani

**Affiliations:** Faculty of Psychology, Department of Clinical and Health Psychology, University of Vienna, Austria; Faculty of Psychology, Department of Cognition, Emotion, and Methods in Psychology, University of Vienna, Austria; Faculty of Psychology, University of Bologna, Italy

**Keywords:** autism spectrum disorder, effective connectivity, functional MRI, interpersonal space, trust game

## Abstract

Previous research indicates that the size of interpersonal space at which the other is perceived as intrusive (permeability) and the ability to adapt interpersonal distance based on contextual factors (flexibility) are altered in Autism Spectrum Disorder (ASD). However, the neurophysiological basis of these alterations remains poorly understood. To fill this gap, we used fMRI and assessed interpersonal space preferences of individuals with ASD before and after engaging in cooperative and non-cooperative social interactions. Compared to matched controls, ASDs showed lower comfort in response to an approaching confederate, indicating preference for larger interpersonal space in autism (altered permeability). This preference was accompanied by reduced activity in bilateral dorsal intraparietal sulcus (dIPS) and left fusiform face area (FFA), regions previously shown to be involved in interpersonal space regulation. Furthermore, we observed differences in effective connectivity among dIPS, FFA, and amygdala in ASDs compared to controls, depending on the level of experienced comfort. No differences between groups were observed in interpersonal space regulation after an experienced social interaction (flexibility). Taken together, the present findings suggest that a dysregulation of the activity and connectivity of brain areas involved in interpersonal space processing may contribute to avoidance of physical proximity and social impairments in ASD.

## INTRODUCTION

In our everyday interactions, we set boundaries to keep others at a preferred distance. The space that we interpose between us and other people is termed “interpersonal” space. It represents a safety zone surrounding the body, evolutionarily developed in order to signal possible social threats approaching us (Graziano and Cooke 2006) and/or to foster feelings of intimacy with conspecifics (Gibson et al. 1993; Sorokowska et al. 2017). An intact capacity to set and regulate interpersonal space preferences is therefore fundamental for health and well-being, as well as for optimal social functioning (Hall 1996).

Two preeminent features need to be taken into account when analyzing interpersonal space differences: *permeability* and *flexibility* (Hayduk 1981). *Permeability* refers to the size of the space at which the others’ approach is perceived as an intrusion and arouses discomfort. *Flexibility* indicates the ability to regulate interpersonal space depending on situational and social factors, such as the level of closeness and familiarity with the other, as well as age, gender, culture, and perceived trustworthiness (Remland et al. 1995; Iachini et al. 2016; Sorokowska et al. 2017; Rosenberger et al. 2020). Notably, both features have been documented to be impaired in Autism Spectrum Disorder (ASD), but with somehow inconsistent results. For example, two previous studies have shown that ASD children have altered *permeability*, as indicated by a preference for larger distances towards an unfamiliar adult (Gessaroli et al. 2013; Candini et al. 2017, 2019), compared to matched typically developing children. However, preference for a smaller interpersonal distance in ASD compared to typically developing children and adolescents has also been reported (Pedersen et al. 1989; Parsons et al. 2004; Asada et al. 2016). In the adult population, Kennedy and Adolphs (2014) found that individuals with ASD intrude more often others personal space than CTRs, while Perry and colleagues (2015) have reported greater variance in interpersonal space preferences in ASD compared to a control sample.

Considering *flexibility*, we have previously documented that typically developing children shrink their interpersonal space following a brief cooperative interaction with a confederate, while they do not modify it after an uncooperative one (Gessaroli et al. 2013; Candini et al. 2017, 2019). In contrast, in ASD children with severe social impairment, no changes in interpersonal space preferences emerged in response to the same cooperative and uncooperative social interactions, suggesting a link between the ability to integrate new information in the regulation of interpersonal space and the impairment in everyday social life (Candini et al. 2017). Notably, to date, no study has investigated changes in interpersonal space flexibility in a population of adults with ASD.

At the neural level, our understanding of interpersonal space processing relies on neuroimaging studies that have used images of virtual human faces or objects presented in different sizes to recreate the effect of individuals approaching towards (or withdrawing from) the participants (Holt et al. 2014; Vieira et al. 2017, 2020). These studies have identified a network of parietal and frontal regions involved in *permeability*. In particular, two key brain areas seem to preferentially respond to social (compared to non-social) stimuli intruding the interpersonal space: the dorsal intraparietal sulcus (dIPS) and the ventral premotor cortex (PMv) (Holt et al. 2014; Vieira et al. 2017, 2020). Notably, activity in these regions has been linked to individuals’ self-reported social activity, such as the amount of time spent, and preferred to spend, with others (Holt et al., 2014). Besides frontal and parietal regions, approaching faces are also processed by the dorsal and ventral visual stream, and have been associated with activity of the fusiform face area (FFA) and the human middle temporal visual area (hMT+/V5) (Holt et al. 2014, 2015; Schienle et al. 2015, 2017). Furthermore, limbic regions, such as the amygdala, have been found to respond to violation of interpersonal space (Kennedy et al. 2009). While hyperactivation of the amygdala has been suggested as a possible mechanism behind the alterations observed in autism (Gessaroli et al. 2013; Candini et al. 2020), to date this hypothesis has not yet been investigated and the neural underpinnings of interpersonal space processing in the ASD population remain largely unknown.

In the present study, we aimed to ascertain the behavioral and neural correlates of interpersonal space processing in adults with ASD. To this aim, we conducted an fMRI study to investigate i) interpersonal space intrusion (*permeability*) and ii) the impact of cooperative and non-cooperative social interactions on interpersonal space preferences (*flexibility*) in a sample of adults with ASD and matched CTRs. A novel ecologically valid task, consisting of videos of real confederates walking towards the participants lying in the scanner, was employed. The task was performed before and after an online “cooperative” and “non-cooperative” social interaction with the two confederates, attained by means of an economic-social game (Rosenberger et al. 2020). Task-based univariate fMRI analyses were performed to assess group differences in the local cortical activity of interpersonal space processing. Dynamic causal modeling (DCM) was implemented to investigate changes in effective connectivity as a possible mechanism, and specifically whether amygdala functions as a node regulating the cross talk between brain regions involved in the processing of interpersonal space.

## MATERIALS AND METHODS

### Sample

20 adults (14 males) with high-functioning ASD and 20 (14 males) controls (CTR) took part in the study (*M*_age_ = 33.11, *SD* = 11.12). We planned to test a minimum sample size of 40 participants (20/group), based on the sample size of previous imaging studies investigating interpersonal space processing in neurotypical and clinical populations (N = 22, Holt et al., 2014; Nstudy1 = 29 and Nstudy2 = 35, Holt et al., 2015; N = 50, Schienle et al. 2015; N = 35, Schienle et al. 2017), and on the availability of adult individuals with ASD in Vienna and surroundings. Participants in the two groups were matched for age, gender, handedness, and intelligence (Table 1), with the latter assessed using the Multiple Choice Vocabulary test (MWT-B, Lehrl et al. 1995) and the Standard Progressive Matrices (SPM; Kratzmeier and Horn 1979). ASD participants had a confirmed clinical diagnosis of ASD according to ICD-10 criteria, provided by an accredited institution and preferentially assessed using the Autism Diagnostic Observation Schedule (Lord et al. 2000). Exclusion criteria for all participants were any contraindication to MRI scanning and studying/having studied psychology.

**Table 1.**
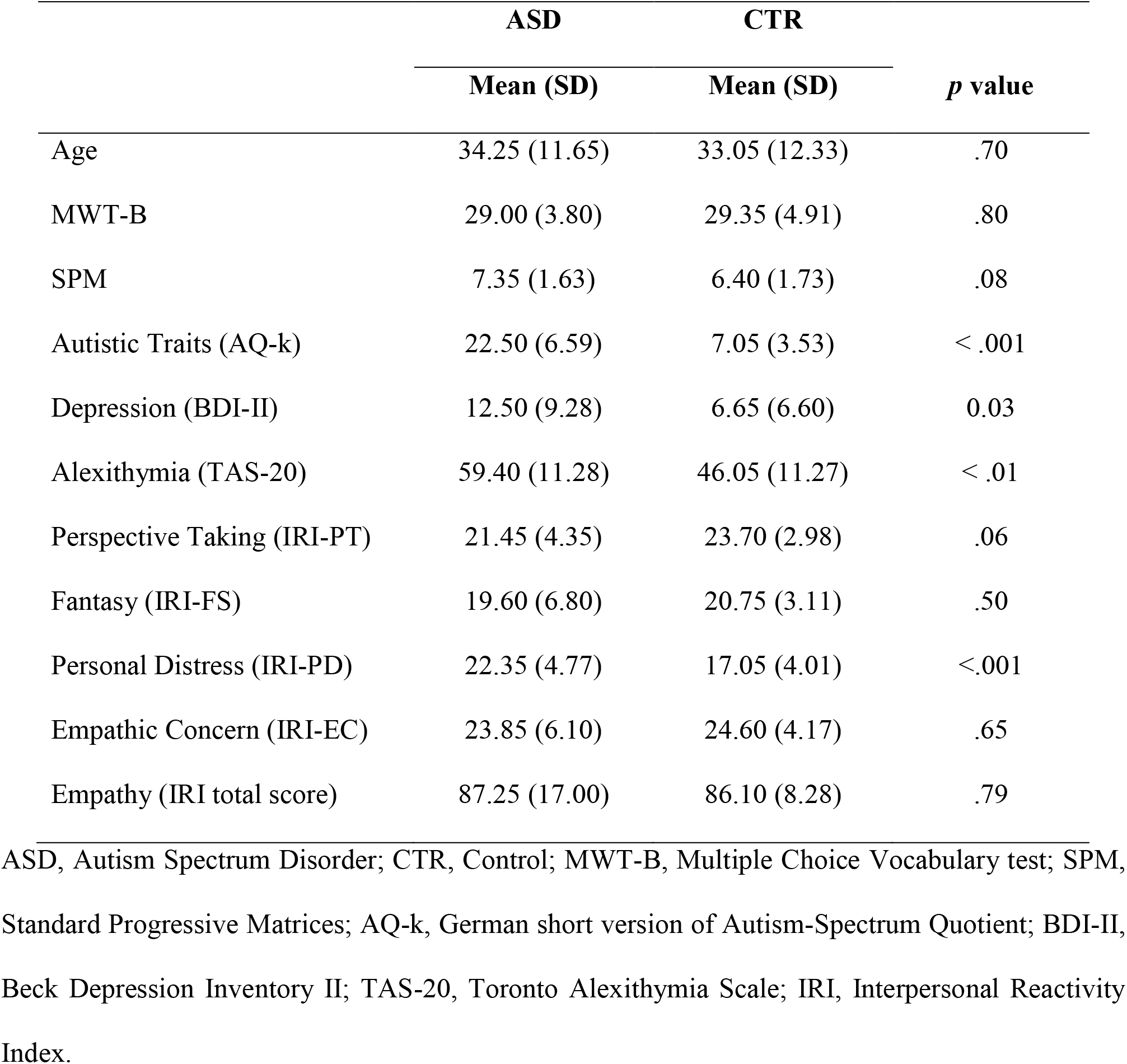
Characteristics of the ASD and CTR samples.

Additional exclusion criteria for controls were psychiatric or neurological disorders and regular medication intake. All participants gave written consent and received monetary compensation of 30 €. The study was approved by the Ethics Committee of the Medical University of Vienna (EK 1166/2015) and complied with the Declaration of Helsinki (World Medical Association 2013).

### Procedure

At the beginning of the experimental session, each participant was introduced to two unknown same-gender confederates, who were presented as two other participants of the study. Participants and confederates were instructed about the experimental tasks and asked to sign the consent form. Participants were then placed into the MRI scanner. In order to investigate how cooperative and non-cooperative social interactions influence interpersonal space regulation, the Interpersonal Space task was performed before (T1) and after (T2) the Repeated Trust Game. In order not to raise any suspicions about the aim of the study, participants were not informed beforehand about the second run of the Interpersonal Space task, but only after the Repeated Trust Game, with the excuse that due to technical problems the task needed to be repeated. At the end of the experimental session, participants filled out self-report questionnaires and were debriefed about the deception.

### Interpersonal Space task

A novel and ecologically valid task was developed and implemented in MatLab R2010a (The MathWorks, Inc., Natick, Massachusetts, United States). Participants were informed that, during the task, an MRI-compatible video camera mounted on the scanner would record and stream online the two confederates walking between one and five steps towards them on a screen behind the scanner, accessible via a mirror system. After observing the confederates approaching them, participants were asked to judge their comfort regarding the distance at which the confederates stopped.

In reality, participants were presented with pre-recorded videos, displaying the confederate walking towards the MRI scanner, at a pre-defined speed (one step per second). A total of 20 videos (one to five steps, for each of the four confederates) were recorded in the room where later the study took place. To reduce variability in appearance, all confederates wore similar clothes and kept the same haircut for the whole duration of the study. During recording, confederates were instructed to keep a neutral facial expression and to direct their gaze on a mark to the right of the camera. A metronome was used to ensure that the confederates walked at the desired speed, while pre-marked locations on the floor of the scanner room were used to ensure that each confederate stopped at the same distance.

The task consisted of a total of 50 trials. For each trial (Fig. 1A), after the fixation cross (5-7s), a video depicting one of the two confederates walking between one and five steps toward the scanner (3-6s) was displayed (Fig. 1B). Participants were then instructed to rate their comfort (“How did you experience the distance to the other person?”) using a Visual Analogue Scale (VAS) ranging from −10 (unpleasant) to +10 (pleasant), with no time limit. Each video was presented five times in random order.

**Fig. 1.**
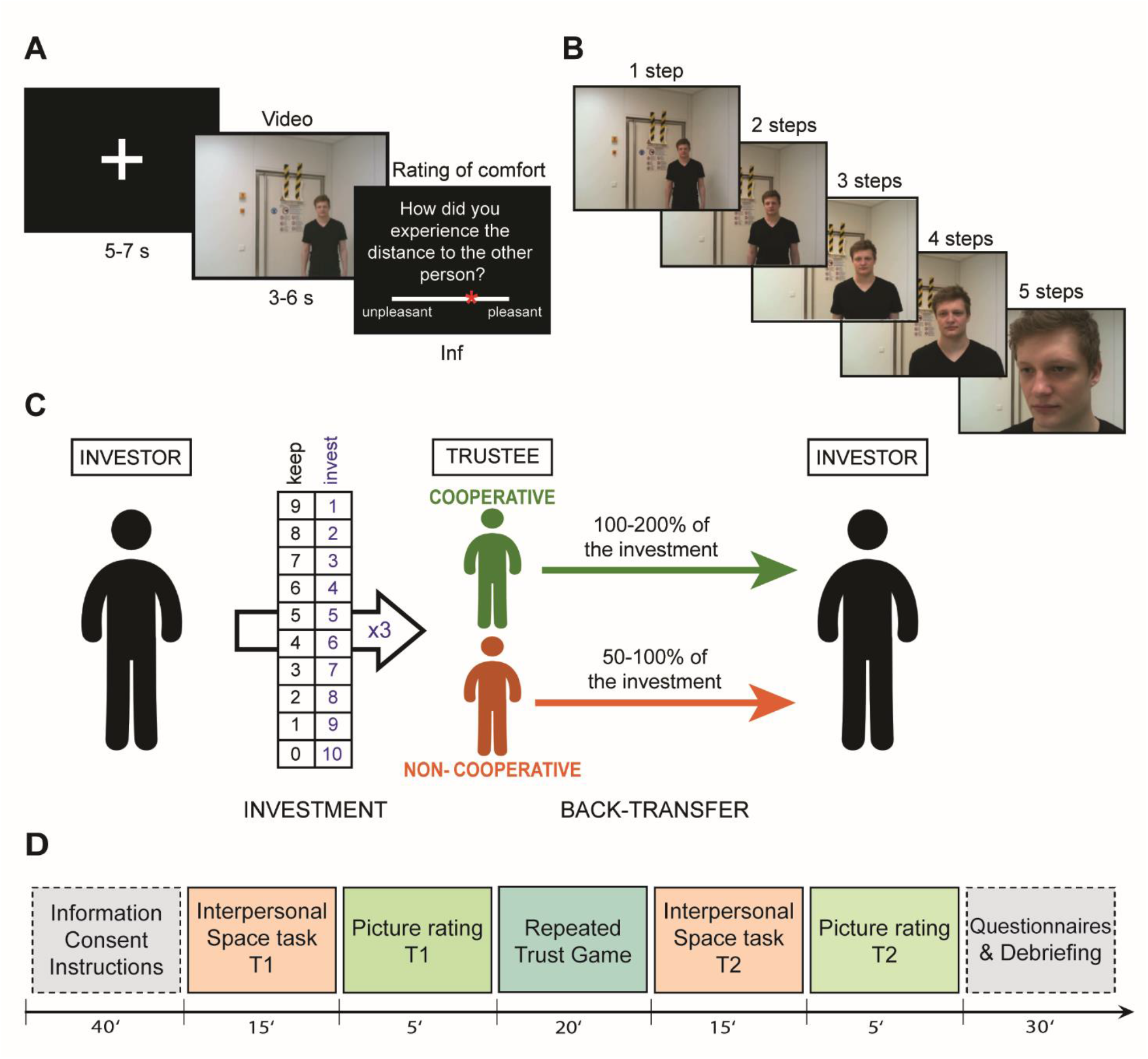
Schematic representation of the experimental tasks and study outline. (A) Trial structure of the Interpersonal Space task. “Inf” indicates that participants could respond with no time limit. (B) Example of last frame of the videos representing the five possible interpersonal distances observed by participants during the Interpersonal Space task (step 1 to 5). (C) Trial structure of the Repeated Trust Game. (D) Overview of the study procedure.

At the end of the task, participants were shown a pre-recorded picture^1^ of each confederate (3s) and asked to rate their perceived fairness, intelligence, attractiveness and trustworthiness on a VAS ranging from −10 (not at all) to +10 (very), with no time limit.

### Repeated Trust Game

The Repeated Trust Game (Berg et al. 1995) was used to manipulate the type of social interaction (cooperative or non-cooperative) experienced by the participants during the study. The trust game is a well-validated economic-social paradigm where two players, an investor and a trustee, exchange money to maximize their profits. In the current task, participants always took the role of the investor and played with the two confederate trustees over multiple trials. Roles were assigned by means of a pre-determined lottery^2^, in which participants were always identified as player A (investor) and the confederates as players B and C (trustees).

The task consisted of 40 trials. Participants played 20 trials with each confederate, in a randomized fashion, while lying in the MRI scanner, with the trustees supposedly connected from laptops placed outside the scanner room. In each trial (Fig. 1C), i) the investor (participant) and the trustee (one of the confederates) received an initial endowment of 10 monetary units (MU) and the picture of the current trial’s trustee was displayed, ii) the investor transferred a part (1 to 10 MU) of their endowment to the trustee (investment), iii) the investment was tripled by the experimenter, thus the trustee received three times the investment (multiplication), iv) the trustee transferred between 1 and 10+3*investment MU back to the investor (back-transfer). The back-transfer behavior of the trustees was computer-controlled so that participants always played with a cooperative (returning between 100 and 200% of the investment) and a non-cooperative (returning between 50 and 100% of the investment) trustee (see Rosenberger et al. 2020 and *Supplementary Material* for a detailed description). The Repeated Trust Game was implemented in Z-tree version 3.4.2 (Fischbacher 2007).

### Questionnaires

At the end of the scanning session, participants filled out a survey regarding socio-demographic information and self-report questionnaires to assess personality traits: the short version of the Autism-Spectrum Quotient (AQ-k; Freitag et al. 2007), the Interpersonal Reactivity Index (IRI; Koller and Lamm 2014), the Beck Depression Inventory (BDI-II; Kühner et al. 2007) and the Toronto Alexithymia Scale (TAS-20; Bach et al. 1996). Lastly, participants were asked to answer some questions to indirectly investigate the credibility of the cover story (see *Supplementary Material*).

### fMRI data acquisition

A 3 Tesla Siemens MAGNETOM Skyra scanner equipped with a 32-channel head coil was used. The scanning sequence parameters of the functional scans were as follows: TE/TR = 34/704 ms, 32 axial slices coplanar with the line connecting anterior and posterior commissure, slice thickness = 3.5 mm, flip angle = 50°, interleaved acquisition, and interleaved multi-slice mode, matrix size = 96×96, voxel size = 2.2×2.2×3.5 mm, field of view = 210 mm. Between ~ 900 and ~1100 volumes were acquired in each run of the Interpersonal Space task and between ~1300 and ~1700 in the Repeated Trust Game^3^.

### Statistical analysis

#### Behavioral data

To investigate differences in *permeability* and *flexibility* of interpersonal space between ASD and CTR groups, we performed a mixed-model Analysis of Variance (ANOVA) on the comfort ratings (Interpersonal Space task) with Group (ASD, CTR) as between-subjects factor and Time (T1, T2), Trustee (Cooperative, Non-cooperative), Step (1-5) as within-subject factors. A mixed-model ANOVA on the investments (Repeated Trust Game), including Group as between-subjects factor and Trustee as within-subject factor, was used to assess whether participants effectively learned about the different intentions of the investors (cooperative vs. non-cooperative). Additionally, we conducted four mixed-model ANOVAs on the ratings of trustworthiness, fairness, attractiveness and intelligence, including Group as between-subjects factor, and Time and Trustee as within-subject factors. Significant interactions were decomposed using post hoc comparisons with Bonferroni correction. Two-tailed *t*-tests were implemented to control for differences between groups regarding age, intelligence and personality traits. Due to missing data, following response box’s failure in recording the answer, the analyses on trustworthiness, fairness, attractiveness, and intelligence were performed on 34 (17 ASD), 32 (17 ASD), 35 (17 ASD) and 35 (18 ASD) participants respectively. All analyses were conducted in R (R Core Team 2019). We used the function aov_car() of the *afex* package for computing ANOVAs, the function emmeans() of the homonymous package for post hoc comparisons, and the package *ggplot2* to create figures. Means and SDs can be found in the *Supplementary Material*.

### fMRI data

Two participants (one ASD) were excluded due to technical problems during image acquisition, resulting in bad image quality. Overall, 19 participants per group were included in the fMRI analyses. Functional data were pre-processed and statistically analyzed using SPM12 (Wellcome Trust Centre for Neuroimaging, www.fil.ion.ucl.ac.uk/spm).

#### Preprocessing

Each functional volume was realigned to the first image, segmented in gray matter, white matter and cerebrospinal fluid tissues, normalized to a template based on the 152 brains from the Montreal Neurological Institute (MNI), and then smoothed by convolution with a 6 mm full width at half maximum (FWHM) Gaussian Kernel. Slice-timing was not applied because not necessary due to the short TR (Sladky et al. 2011). Motion related parameters were visually inspected for head excessive movements (> 3 mm/degrees) and corrected by replacing the problematic images with a mean image calculated from the preceding and succeeding images.

#### Task-based univariate fMRI analysis

Statistical analysis was performed using a general linear model approach (Friston et al. 1994), involving a two-step procedure (first and second level analyses). In the first-level analysis, regressors of interest were defined for each participant and convolved with the canonical hemodynamic response function. High-pass temporal filtering with a cut-off of 128 s was used to remove low-frequency drifts. For each run, 11 regressors were defined, one for each video type (Cooperative, Non-Cooperative confederate and Step 1 to 5) plus one for the ratings. The inter-trial intervals (fixation cross) served as an implicit baseline. Residual effects of head motion were corrected by including the estimated motion parameters of each participant as six additional regressors of no interest in the design matrix. For the second-level analysis, first-level contrast images derived from the simple effect of each video type were fed into a full-factorial ANOVA design with the between-subjects factor Group (ASD, CTR) and three within-subject factors Time (T1, T2), Trustee (Cooperative, Non-Cooperative) and Step (1-5), using a random-effects analysis (Holmes and Friston 1998). Linear contrasts of this ANOVA model were used to assess main effects and interactions. In order to investigate differences between factors in regions previously shown to be responsive to interpersonal space intrusion, a mask including bilateral IPS, PMv, amygdala, FFA, and hMT+/V5 was used for small volume corrections (SVC). The mask was built by combining spheres of 8 mm centered on the peak coordinates reported in previous papers on interpersonal space intrusion (FFA and hMT+/V5: Foss-Feig et al., 2016; IPS and PMv: Holt et al., 2014, 2015; amygdala: Kennedy et al., 2009), using the MarsBaR SPM toolbox (http://marsbar.sourceforge.net/; see *Supplementary Material* for a list of the peak coordinates used). The reported results are based on family-wise error (FWE) correction for voxel intensity tests (*P*_FWE_ < .05).

### Correlation analysis

Pearson correlations with Bonferroni correction were performed to explore the associations between neural activity and a) averaged comfort ratings expressed by the participant during the Interpersonal Space task, and b) reported social abilities, as indicated by the score of the subscale “Social interaction and spontaneity” of the AQ-k. We focused on the social subscale of the AQ-k based on evidence reported for a positive correlation between dIPS and PMv, and participants’ level of sociability (Holt et al. 2014). Spheres of 4 mm centered on the peak voxels resulted from the main contrast CTR>ASD were built using MarsBar SPM toolbox. Activity extracted from those spheres was then correlated with the individual scores.

### Dynamic causal modelling (DCM)

Previous literature hypothesized the amygdala as a possible key structure at the base of the impairments in interpersonal space regulation in ASD. Therefore, we explored whether the connectivity between areas showed to be differently activated in ASDs and CTRs in the task-based univariate analysis, i.e. FFA and dIPS, and the amygdala was altered in ASD compared to CTR. A two-steps procedure was implemented, consisting of a first-level DCM analysis (Friston et al. 2003) performed for each subject, followed by group-level modelling using the Parametric Empirical Bayes framework (Friston et al. 2016). Group and comfort ratings were entered as covariates of interest in the second/group-level analysis.

More precisely, BOLD time series (principal eigenvariate) from the masks of dIPS, FFA, and amygdala used for SVC in the GLM analysis, and from anatomical mask of V1 (hO1) included in SPM Anatomy Toolbox (www.fz-juelich.de), were extracted from the main effect of video’s presentation for each participant^4^. Since a group difference in the activity of FFA was found only in the left hemisphere, we focused on the interregional coupling of this hemisphere. Each timeseries then entered a first-level DCM analysis, in which the effect of the videos (vs. baseline) on the reciprocal coupling between amygdala, FFA and dIPS, with V1 as input region, was assessed. The DCM model was estimated for each subject on the first run^5^ of the Interpersonal Space task. Parametric Empirical Bayes (PEB) was implemented. The group-level PEB design matrix included the zero-mean centered covariate of interest Group, the mean-centered covariate Comfort as well as their interaction. The analysis focused on the modulation of the effective connectivity by the task (*B* matrix). In addition, intrinsic connectivity (*A* matrix) was calculated in order to interpret the directionality of the findings of the *B* matrix. Bayesian model reduction (Friston et al. 2016) was used to prune parameters based on the free energy^6^ (Zeidman et al. 2019). We only reported inter-regional

## Data availability

Behavioral data and analysis scripts are available online (https://osf.io/t6sxf/). Unthresholded statistical maps and the used ROI mask are available on NeuroVault (https://neurovault.org/collections/8941/).

## RESULTS

### Behavioral data

#### Interpersonal Space task

The ANOVA conducted to examine differences in comfort (*permeability* and *flexibility* of interpersonal space) between ASD and CTR revealed a significant main effect of Group (*F*_1,38_ = 5.66, *p* = .023, η^2^_G_ = .04), Trustee (*F*_1,38_ = 11.51, *p* = .002, η^2^_G_ = .010), Time (*F*_1,38_ = 4.69, *p* = .037, η^2^_G_ = .002), and Step (*F*_1.7,63.5_ = 67.64, *p* < .001, η^2^_G_ = .517). Regarding the main effect of Group, participants in the ASD group showed a general lower comfort (i.e., reduced *permeability*) while watching the confederates approaching, compared to the CTR group (Fig. 2A). As expected, participants’ comfort generally decreased as the distance between them and the confederate became smaller^7^. Additionally, the analysis revealed the following significant interactions: Time x Trustee (*F*_1,38_ = 19.99, *p* < .001, η^2^_G_ = .006), Time x Step (*F*_3.2,121.6_ =8.20, *p* < .001, η^2^_G_ = .006) and Time x Trustee x Step (*F*_2.8,106.4_ = 3.76, *p* = .015, η^2^_G_ = .002). Decomposing the Time x Trustee x Step interaction (Fig. 2B), post hoc comparisons revealed that, after the Repeated Trust Game, when watching the cooperative confederate approaching them, participants increased their comfort ratings at step 4 (*p* < .001). On the other hand, when watching the non-cooperative confederate approaching them, participants decreased their comfort ratings, in particular at larger distances (step 2 and 3, all *p* < .001). Contrary to our hypothesis, we did not find any significant interaction with the factor Group, meaning that the modulation of interpersonal space preferences by a positive or negative social interaction (i.e., *flexibility)* was similarly present in ASD and CTR participants.

**Fig. 2.**
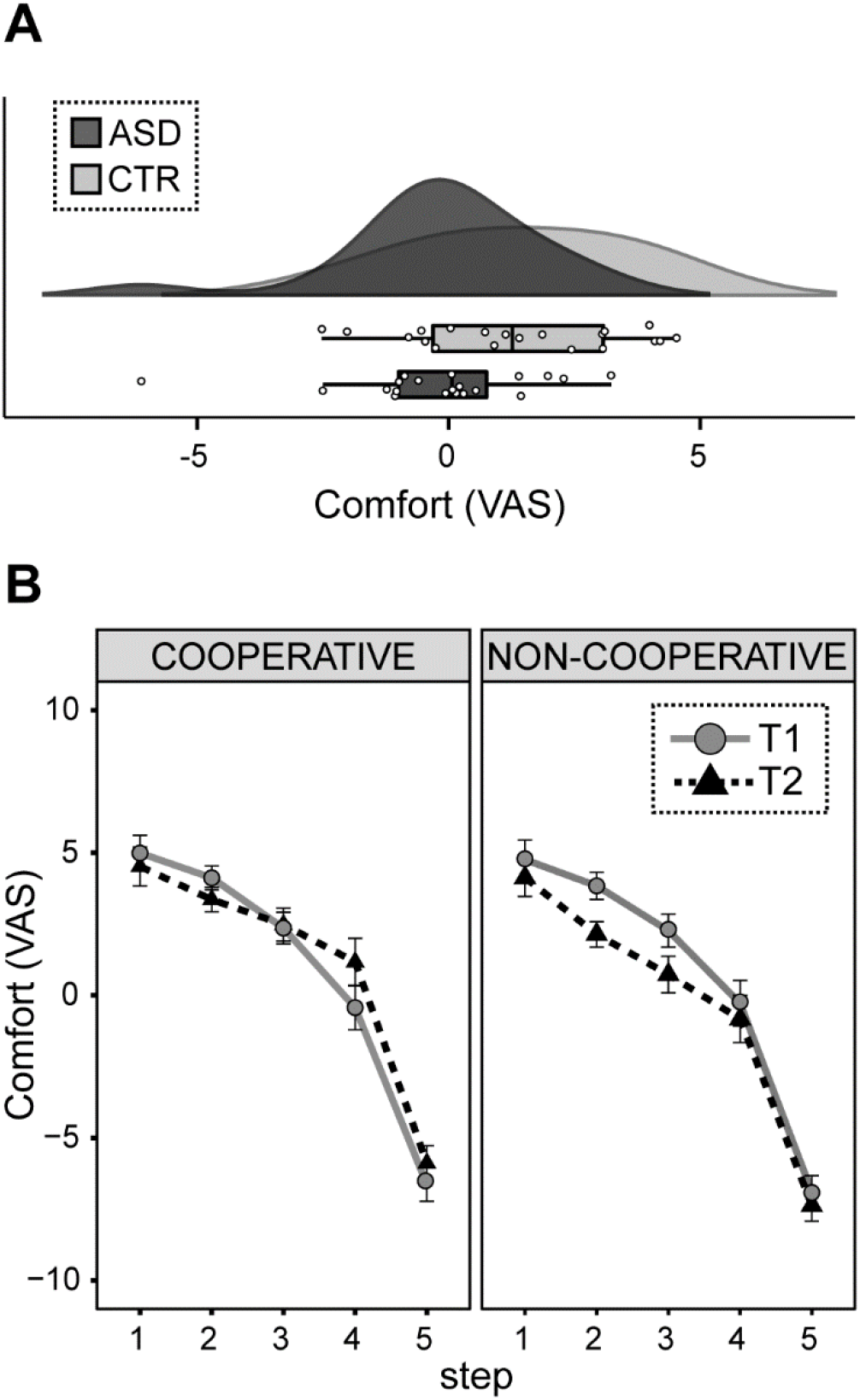
Behavioral results of the Interpersonal Space task. (A) Raincloud plot of the averaged comfort ratings expressed by the ASD and the CTR group during the Interpersonal Space task. The half violin plots depict the probability density of the data at different values. Dots represent the means of individual subjects. (B) Comfort ratings expressed for the cooperative and non-cooperative trustee for the different distances (step 1 to 5). T1, run of the Interpersonal Space task performed before the Repeated Trust game; T2, run of the Interpersonal Space task performed after the Repeated Trust game. Error bars represent the standard error of the mean. CTR, control; ASD, Autism Spectrum Disorder.

#### Repeated Trust Game

The ANOVA conducted on participants’ investments showed a significant main effect of Trustee (*F*_1,38_ = 178.61,*p* < .001, η^2^_G_ = .62), indicating that participants from both groups learned to maximize their profit during the task by investing more in the cooperative trustee and less in the non-cooperative one (Fig. 3A). The absence of a significant main effect or interaction with the factor Group (all *F*< 3.93, all *p* > .19) confirmed that this learning was similar for both groups.

**Fig. 3.**
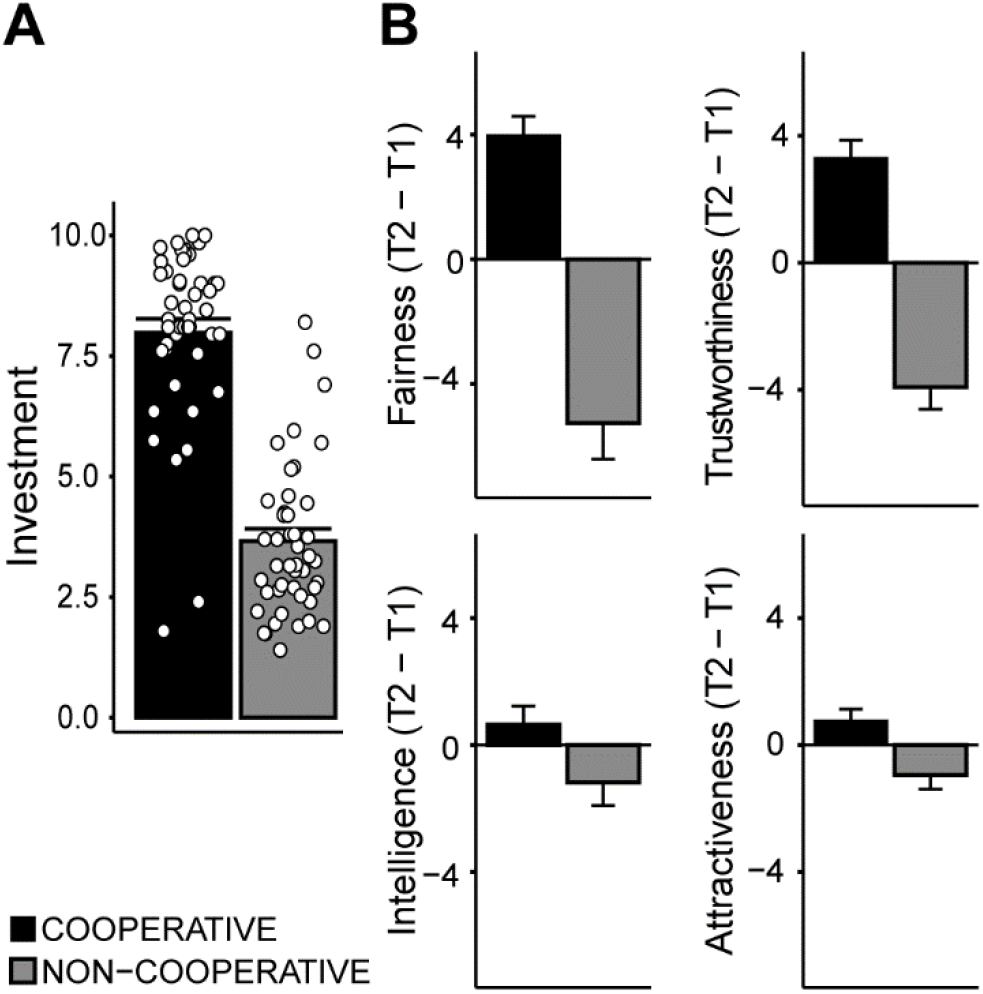
Behavioral results of the Repeated Trust Game and picture rating. (A) Mean investments in MU for the cooperative and non-cooperative trustee in the Repeated Trust Game. (B) Changes (T2 - T1) in the ratings of fairness, trustworthiness, attractiveness, and intelligence after cooperative and non-cooperative social interactions. Positive values reflect an increase while negative values reflect a decrease in the judgement expressed in the second run (after the Repeated Trust Game) compare to the first run. T1, run of the Interpersonal Space task performed before the Repeated Trust game; T2, run of the Interpersonal Space task performed after the Repeated Trust game. Dots represent the means of individual subjects. Bars represent standard error of the mean.

In agreement with these findings, the ANOVAs on trustworthiness and fairness ratings revealed a significant main effect of Trustee (trustworthiness: *F*_1,32_ = 12.25, *p* < .01, η^2^_G_ = .14; fairness: *F*_1,30_ = 17.89, *p* < .001, η^2^_G_ = .16) and a significant Time x Trustee interaction (trustworthiness: *F*_1,32_ = 50.26,*p* < .001, η^2^_G_ = .26; fairness: *F*_1,30_ = 31.71,*p* < .001, η^2^_G_ = .22). Post hoc comparisons showed that, after the Repeated Trust Game, participants significantly increased and decreased their judgment of trustworthiness and fairness of the cooperative and non-cooperative confederate respectively (all *p* < .001; Fig. 3B). Similarly, the ANOVA on intelligence ratings revealed a trend to significance for the main effect of Trustee (*F*_1,33_ = 3.42, *p* = .07, η^2^_G_ = .03) and for the Time x Trustee interaction (*F*_1,33_ = 3.01, *p* < .09, η^2^_G_ = .01; Fig. 3B). Lastly, the ANOVA on attractiveness ratings revealed a trend to significance for the main effect of Trustee (*F*_1,33_ = 3.16, *p* = .08, η^2^_G_ = .02) and a significant Time x Trustee interaction (*F*_1,33_ = 7.03, *p* < .05, η^2^_G_ = .01; Fig. 3B). However, post hoc comparisons of attractiveness before and after the social interaction did not reveal significant difference.

Overall, the data indicate that the Repeated Trust Game was effective in inducing a shift in the judgments about the personality of the confederates and that those changes were similar for the ASD and CTR groups.

### Neuroimaging data

#### Permeability of interpersonal space

To assess differences in brain activity related to interpersonal space *permeability* between the ASD and CTR groups, we computed the contrasts CTR > ASD and ASD > CTR for the main effect of videos across the two runs of the Interpersonal Space task.

Results from the contrast CTR > ASD showed significant activation of the left dIPS, right dIPS, right hMT+/V5, and left FFA (Fig. 4A, Table 2). The opposite contrast (ASD > CTR) showed significant activation of the left hMT+/V5.

**Fig. 4.**
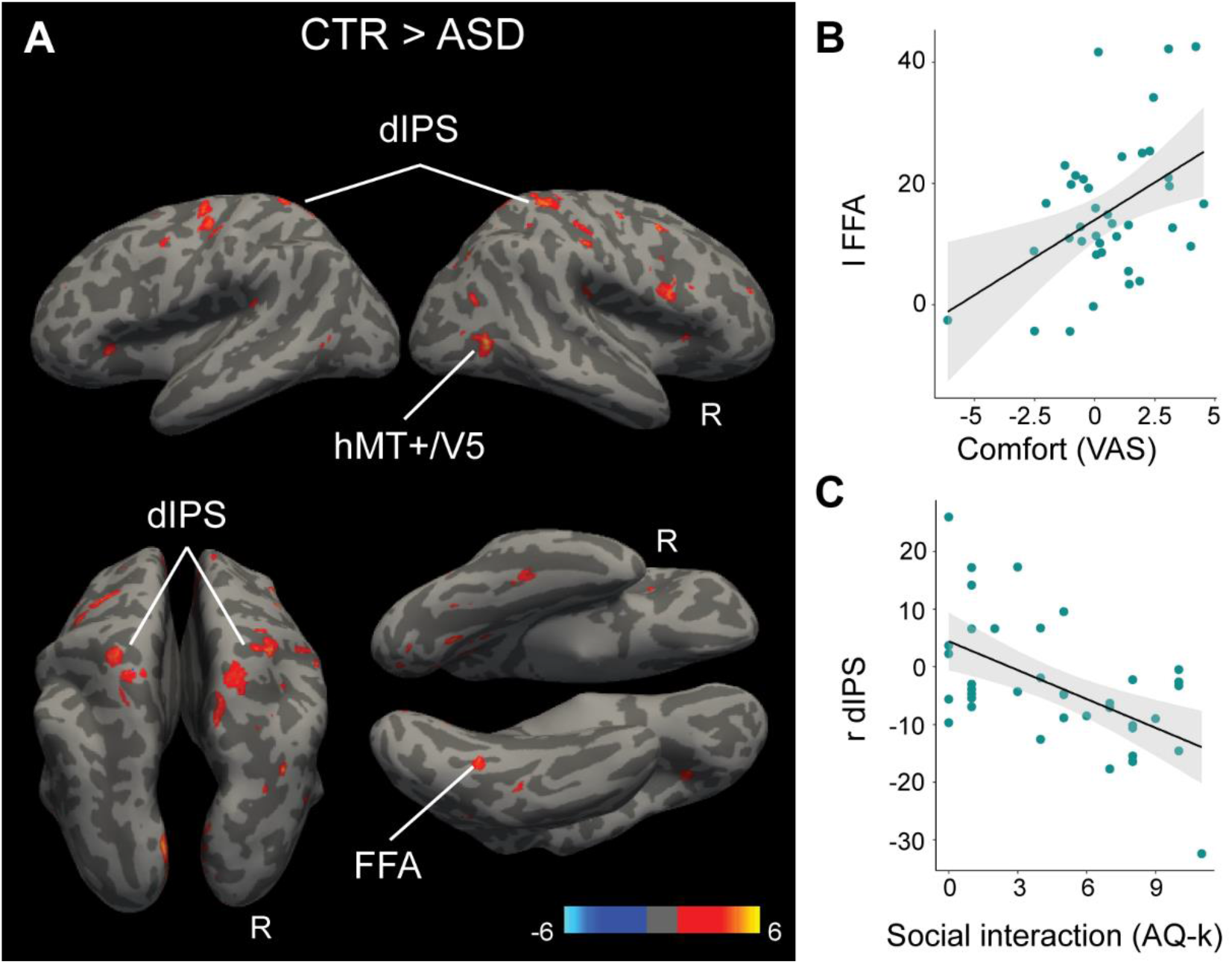
Functional MRI responses during the Interpersonal Space task. (A) Enhanced neural responses to the approaching confederate in bilateral dIPS, right hMT+/V5 and left FFA in CTRs compared to ASDs. (B) Positive correlation between left FFA activity and the averaged comfort expressed during the Interpersonal Space task across the whole sample. (C) Negative correlation between right dIPS activity and the score of the social interaction subscale of the AQ-k questionnaire (higher scores indicate lower social abilities). CTR, control; ASD, Autism Spectrum Disorder; dIPS, dorsal Intraparietal Sulcus; hMT+/V5, human middle temporal visual area; FFA, Fusiform Face Area. Statistical maps are displayed at *p*<.001 uncorrected for visualization purposes.

**Table 2.** Permeability of the interpersonal space

| Region | Hemi | Cluster K | Z | $p_{FWE}$ | x | y | z |
| --- | --- | --- | --- | --- | --- | --- | --- |
| *CTR > ASD* |  |  |  |  |  |  |  |
| Superior Parietal Gyrus<sup>a</sup> | R | 23 | 4.6 | .003 | 30 | -36 | 58 |
| Superior Parietal Gyrus<sup>a</sup> | L | 82 | 4.6 | .004 | -20 | -48 | 62 |
| Inferior Occipital Gyrus<sup>e</sup> | R | 15 | 4.4 | .008 | 42 | -62 | 2 |
| Fusiform Gyrus<sup>d</sup> | L | 38 | 4.1 | .033 | -34 | -54 | -16 |
| *ASD > CTR* |  |  |  |  |  |  |  |
| Middle Occipital Gyrus<sup>e</sup> | L | 21 | 4.5 | .005 | -42 | -76 | 4 |

Voxels which showed significantly ( $p < .05$ FWE-corrected) greater activation in the contrasts of interest. Voxels falling within the predefined ROI mask for dIPS, PMv, amygdala, FFA and hMT+/V5 are indicated with *a*, *b*, *c*, *d* or *e* (*a*, dIPS; *b*, PMv; *c*, Amygdala; *d*, FFA; *e*, hMT+/V5). Voxels were labelled based on John Hopkins Uni atlas (Faria et al. 2012) in MRIcron (www.mricron.com). CTR, control; ASD, Autism Spectrum Disorder.

#### Flexibility of interpersonal space

In order to assess differences in interpersonal space *flexibility* between ASDs and CTRs, we performed linear contrasts examining the effect of Group x Time (T1, T2) x Trustee (Cooperative, Non-cooperative) and Group x Time x Trustee x Step (1-5 steps), on the video segments. None of these contrasts resulted in any significant voxel. We thus conducted an exploratory analysis only in the second run of the Interpersonal Space task (T2), examining the effect of Group x Trustee and Group x Trustee x Step. No significant activations were found.

Lastly, we explored the Time x Trustee and Time x Trustee x Step interactions across the two groups, but did not observed any significant activations.

#### Correlations between questionnaires, behavioral and MRI data

Correlation analyses revealed a significant positive correlation between the activity of left FFA and the averaged comfort ratings expressed by the participants during the task (*r* = .455, *p* = .032), (Fig. 4B), as well as a significant negative correlation between the activity of right dIPS and the score of the “Social interaction and spontaneity” subscale (AQ-k; *r* = −.547, *p* < .001) (Fig. 4C). Note that higher scores in the AQ-k social subscale indicate lower social competences.

#### Dynamic causal modelling (DCM)

We used DCM to explore changes in effective connectivity between left dIPS, FFA, and amygdala during the Interpersonal Space task as a function of Group, Comfort, and their interaction. Regarding the main effect of Group, the ASD group showed increased connectivity from amygdala to dIPS (1.75) and FFA (1.37), as well as reduced connectivity from dIPS to FFA (−0.61) compared to the CTR group (Free energy, all Pp > .99, Fig. 5A). The main effect of Comfort showed that as comfort decreased, connectivity from FFA to amygdala increased (0.27; Free energy, Pp > .99; Fig. 5B). The Group x Comfort interaction revealed that, as comfort decreased, ASDs showed increased connectivity from dIPS to amygdala (0.32; Fig. 5C,D) and reduced connectivity from FFA to amygdala (−0.36; Fig. 5C,E) compared to CTRs (Free energy, all Pp > .99). A table including the complete *A* (instrinsic connectivity) and *C* (direct input) matrices can be found in the *Supplementary Material* (Table S5).

**Fig. 5.**
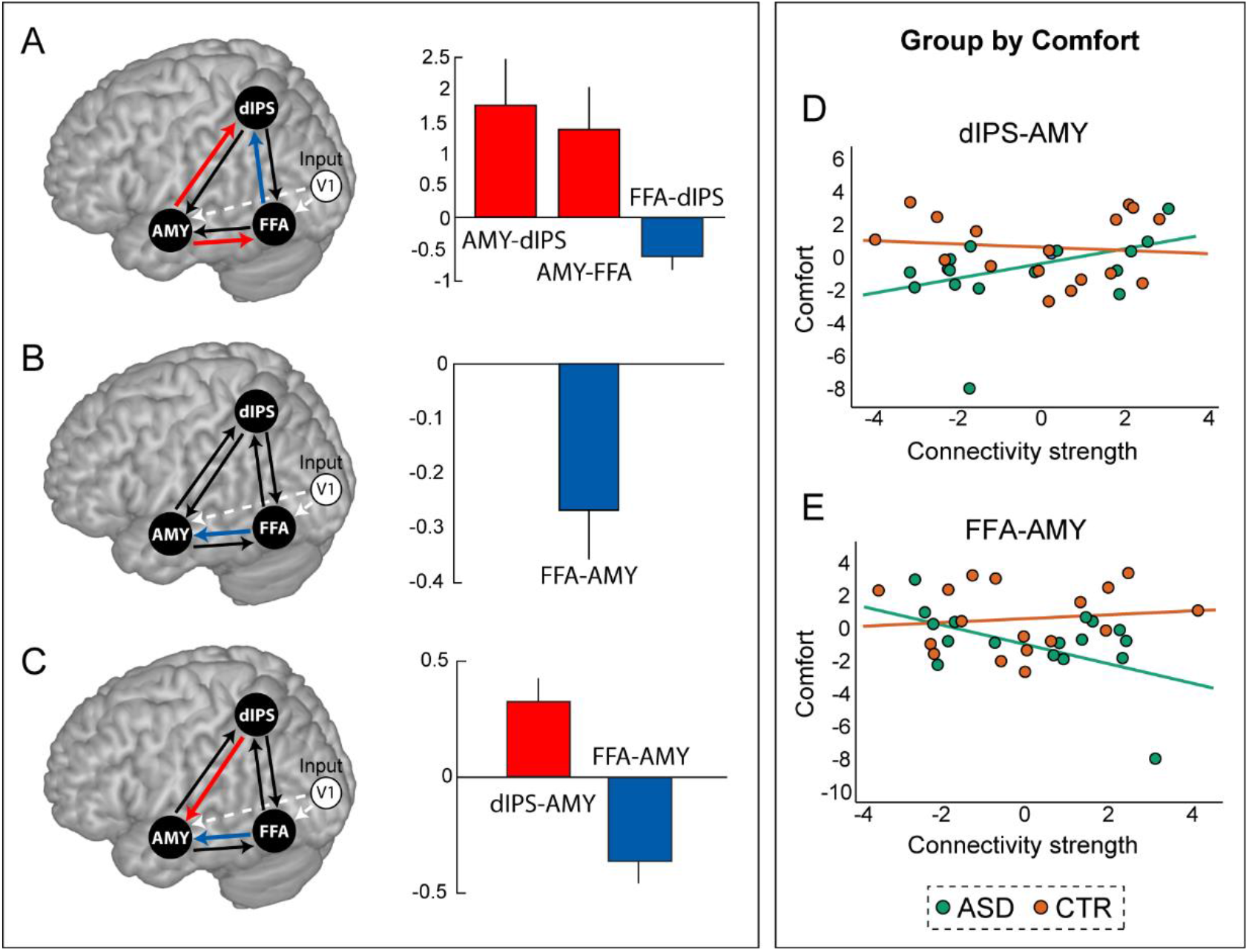
Dynamic causal modelling (DCM). Posterior parameter estimates for the main effects of Group (A) and Comfort (B), and the interaction effect Group x Comfort (C). On the left, red and blue arrows indicate connection parameters exhibiting posterior probability (Pp) > 99%. On the right, bar plots of parameters exhibiting Pp > 99% are displayed. Relationship between averaged comfort expressed during the Interpersonal Space task and dIPS to Amygdala connectivity (D), and FFA to Amygdala connectivity (E) for the two groups (ASD, CTR). Connectivity strengths are shown as unitless log-scaling parameters. CTR, control; ASD, Autism Spectrum Disorder; Amy, Amygdala; dIPS, dorsal Intraparietal Sulcus; FFA, Fusiform Face Area.

## DISCUSSION

The current study examined the behavioral and neural correlates of interpersonal space *permeability* and *flexibility* in adults with ASD and age-, gender-, and intelligence-matched CTRs. Using a new MR-compatible task, we provided evidence that adults with ASD report stronger feelings of discomfort (altered *permeability*), compared to CTRs, when observing another individual approaching them. Furthermore, reduced activity in parietal and visual regions was observed in ASDs compared to CTRs, pointing to a dysregulation of the brain network involved in interpersonal space processing. Using DCM, we identified a possible *neural mechanism* behind such dysregulation, based on different effective connectivity among dIPS, FFA, and amygdala in the ASD group. Finally, we found no evidence for altered *flexibility* of interpersonal space in ASD individuals, compared to CTRs.

### Altered permeability of interpersonal space in ASD

Our results replicate and extend to the adult population the observation of altered interpersonal space *permeability* in individuals with ASD. Previous studies suggested that children with ASD have preferences for larger interpersonal distance (Gessaroli et al. 2013; Candini et al. 2017, 2019; but see also Pedersen et al. 1989; Parsons et al. 2004; Asada et al. 2016) compared to typically developing children. Consistently, our results showed lower comfort in adults with ASD when observing somebody approaching them, indicating an extended interpersonal space. As previously suggested (Candini et al. 2017), ASDs’ impaired social functioning might result in avoiding physical proximity, leading to a decreased experienced comfort when others are approaching.

At the neural level, the altered *permeability* of interpersonal space processing in ASDs was accompanied by reduced activity in the bilateral dIPS, right hMT+/V5 and left FFA. Previous studies showed that dIPS is preferentially activated by approaching social stimuli, as compared to non-social objects, forming, together with PMv and other parietal regions, a fronto-parietal network crucially involved in interpersonal space regulation (Holt et al. 2014; Vieira et al. 2017, 2020). Moreover, an approach-bias for social stimuli in dIPS (i.e., stronger activation for social vs. non-social stimuli) has been found to be positively correlated to individuals’ personal space size and to the amount of time the participants spent, and preferred to spend, with others (Holt et al. 2014). This suggests that the strength of this bias might depend on the amount of experienced social interactions, and thus physical proximity, with others. ASD is a disorder characterized by impairments in the social sphere, where the social network size and amount of social interactions are usually reduced. Therefore, it is possible to hypothesize that the reduced activation of dIPS while observing others approaching may reflect a lower exposure and experience of social proximity in everyday life. In line with this interpretation, we observed a significant negative correlation between right dIPS activity and participants’ level of social ability, as well as a significant positive correlation between left FFA activity and the averaged comfort expressed during the task.

Hypoactivation of FFA during face processing has, since long, been proposed as one of the neuroendophenotypes of ASD (Schultz 2005; Nickl-Jockschat et al. 2015). Notably, previous research has shown that, during the processing of approaching faces, dIPS is functionally connected with areas of the ventral visual stream, in particular the face-responding fusiform area (Holt et al. 2014, 2015). Regions such as FFA and hMT+/V5 provide relevant information to parietal regions about moving social stimuli which are then integrated with information regarding their spatial location, allowing the processing of complex dynamic social stimuli. Dysregulation in this network might therefore result in an impaired integration of the information from the dorsal and the ventral stream, leading to changes in behavior during social interactions, such as the here investigated interpersonal space regulation. By using DCM we were able to show that such impaired integration is due to altered connectivity between amygdala, FFA, and dIPS in the ASD population when processing approaching social stimuli.

### Altered connectivity between amygdala, FFA and dIPS as a possible neural mechanism behind reduced permeability in ASD

Using DCM, we showed differences in effective connectivity between dIPS, FFA, and amygdala between ASDs and CTRs when observing approaching confederates. In particular, we found reduced connectivity from FFA to dIPS in ASDs compared to CTRs, indicating a disruption in the flow of information from the social visual streams to higher order areas that process location during social interaction.

Notably, despite an absence of a group difference in amygdala activity during the observation of approaching individuals, we found increased connectivity from amygdala towards dIPS and FFA in ASD compared to CTRs. The location of potential threats in the environment is an important ability for survival and requires the integration of spatially and motivationally relevant information about an external event. It has been shown that information regarding the motivational relevance from the amygdala and spatial coordinates of an external stimulus are integrated in the spatial attentional network, including dIPS, to form a salience map guiding attention (Egner et al. 2008). Moreover, evidence for greater functional coupling for emotional cues (e.g., angry faces) between amygdala and spatial regions, as well as the fusiform gyrus, suggests that the expectation of threatening stimuli is encoded in the amygdala, which in turn modulates activity in the spatial attention network and infero-temporal visual areas, including FFA, to facilitate the rapid detection of relevant events (Mohanty et al. 2009). Therefore, the increased connectivity from amygdala to dIPS and FFA in ASDs might reflect a higher perceived threat, and thus a higher saliency, of the person approaching. This interpretation is corroborated by the observation that comfort modulates the strength of the connection between FFA and amygdala, with increasing decoupling between those regions when the reported overall comfort was higher. Notably, when we analyzed the interaction between comfort and group, we observed that in ASDs higher discomfort was associated with higher connectivity from FFA to amygdala, and lower connectivity from dIPS to amygdala, as compared to CTRs, where this pattern was reversed (i.e., lower connectivity from FFA to amygdala, and higher connectivity from dIPS to amygdala with higher discomfort). These results seem to suggest that, while normally a threat approaching us leads to increased coupling between spatial, rather than visual, and salience nodes (i.e., it is more important to track the *where* rather than the *what*), in the case of autism this coupling is functioning differently. Such alteration can possibly give rise to discomfort and the necessity to compensate by keeping the other at a larger interpersonal distance.

### Preserved flexibility of interpersonal space in ASD

Concerning *flexibility* of interpersonal space, the effectiveness of the manipulation of the nature of the social interaction was confirmed in both experimental groups. Indeed, after the Repeated Trust Game, confederates behaving in a cooperative manner were perceived as more trustworthy and fair, while the opposite pattern was found for the confederates acting in a non-cooperative way.

Importantly, we observed that engaging in a cooperative interaction led to a shrinking of the interpersonal space, while engaging in an uncooperative interaction caused the interpersonal space to expand. This was indicated by the fact that, after the Repeated Trust Game, participants increased their comfort when seeing the cooperative confederate approaching, especially when they were stopping at a very short distance from the participant (step four). On the other hand, participants decreased their comfort when seeing the uncooperative confederate approaching, especially when they were stopping at an intermediate distance from the participant (steps two and three). Notably, no evidence was found in support of an impairment in the ability to flexibly adjust interpersonal space based on the nature of the experienced social interaction in adults with ASD. Our clinical sample was constituted of high-functioning individuals, while previous findings suggested that only autistic children with severe impairments in the social sphere present difficulties in flexibly adjusting the interpersonal space based on social factors (Candini et al. 2017). Future studies investigating autistic adults with high vs. low social impairment will be needed to explore whether this evidence replicates in the adult population. Another possible explanation is that the manipulation of social interaction used in previous studies might have failed in inducing comparable effects in CTRs and ASDs. Therefore, the observed differences in the ability to flexibly regulate interpersonal space based on the experienced social interaction between the two groups might have been the result of a sub-optimal manipulation rather than a lack of modulation of interpersonal space. In the current study, the similarities in the investments towards the cooperative and non-cooperative confederate, as well as the resulting shift in judgments of trustworthiness and fairness, indicate that both groups were able to identify the different nature of the social interaction with the confederates and to consequently change their perception of the individual. Nevertheless, no clear differential pattern of brain activity was observed before and after the social interaction across both groups. Future studies are needed to further investigate whether and how this effect is represented at the neural level.

## Conclusion

In conclusion, this study provides evidence for an altered *permeability* of interpersonal space in adults with ASD. Participants with ASD showed higher level of discomfort during the processing of approaching individuals, associated with decreased activity in parietal and infero-temporal regions and altered connectivity between those regions and the amygdala. A dysregulation of the interpersonal space brain network and the associated increased discomfort for approaching others, possibly due to reduced salience of spatial information (i.e. where) compared to visual information (i.e. what), might contribute to the avoidance of physical proximity and the impaired social abilities characterizing autism. Interventions aimed at restoring the salience of spatial information may help ameliorating interpersonal space regulation, and possibly the general social functioning of individuals with ASD. Finally, it is important to note that our conclusions refer to high functioning adult individuals. Future studies will be needed to extend the investigation of interpersonal space processing also to low functioning adults, largely underrepresented in neuroimaging research on ASD (Jack and Pelphrey 2017). This will require the use of simplified fMRI compatible paradigms that enhance compliance with the experimental tasks, by reducing for instance the time spent in the scanner.

## AKNOWLEDGMENTS

We thank Ronald Sladky for his support concerning the DCM analysis and for commenting on a previous version of the manuscript. We further thank Mareike Hubinger and Sabrina Scala for their contribution to task preparation and data collection, as well as the confederates Silvia Deneva, Lukas Kraiger, Marie Pellegrini, and Patrick Smela.

## AUTHORS CONTRIBUTIONS

GS, GDP, FF, MC conceived and planned the study. CM, AG and HH collected the data. CM analyzed the data and wrote the first draft of the manuscript under the guidance of GS, GDP, FF and MC. LR designed and programmed the Repeated Trust Game. All authors contributed to and approved the final version of the manuscript.

## Abbreviations

ANOVA: Analysis of Variance
ASD: Autism Spectrum Disorder
AQ-k: German short version of Autism-Spectrum Quotient
BDI-II: Beck Depression Inventory II
CTR: Control
DCM: Dynamic Causal Modeling
FFA: Fusiform Face Area
dIPS: dorsal Intraparietal Sulcus
IRI: Interpersonal Reactivity Index
hMT+/V5: human Middle Temporal visual area
MWT-B: Multiple Choice Vocabulary test
PEB: Parametric Empirical Bayes
PMv: ventral Premotor Cortex
Pp: Posterior probability
SPM: Standard Progressive Matrices
TAS-20: Toronto Alexithymia Scale
VAS: Visual Analog Scale

## Footnotes

1 In order to enhance the credibility of the procedure, pictures of the participant and confederates were actually taken at the beginning of the experimental session by the experimenter but later not used for the task.

2 Participants were led to believe that this was a real lottery in which the role of investor and trustee were randomly assigned to the three participants.

3 The volumes acquired during the Repeated Trust Game will be pre-processed and analysed separately and not further discussed in the current paper.

4 Note that we did not include hMT+/V5 in the model for two main reasons: 1) in order to simplify the model and 2) because both high and low activity of this region was observed for the Group comparison.

5 Since concatenation of experimental runs with considerable time interval in between is problematic for DCM calculation, we decided to focus only on the first Interpersonal Space task run.

6 The free energy is the sum of all subjects’ DCMs accuracies, minus the complexity induced by fitting the DCMs and the second-level GLM connections which survived the variational free energy based threshold of posterior probability (Pp) greater than 0.99.

7 Notably, the same significant main effects of Group and Step were detected when considering only the Interpersonal Space task before the social interactions (T1, see *Supplementary Material* for full details), leading us to assume that the findings were not dependent on the experimental manipulation of social interaction.

