## Supplementary Material for "Neural correlates of interpersonal space permeability and flexibility in autism spectrum disorder"


Claudia Massaccesi [1],

Alexander Groessing [1],

Lisa A. Rosenberger [2],

Helena Hartmann [2],

Michela Candini [3],

Giuseppe di Pellegrino [3],

Francesca Frassinetti [3],

Giorgia Silani [1*]

[1] Faculty of Psychology, Department of Clinical and Health Psychology, University of Vienna, Austria

[2] Faculty of Psychology, Department of Cognition, Emotion, and Methods in Psychology, University of Vienna, Austria

[3] Faculty of Psychology, University of Bologna, Italy

Index

1. Detailed description of the Repeated Trust Game
2. Validity of the cover story
3. Results from ANOVA performed at T1 (behavioral data)
4. Supplementary tables
    a. Table S1
    b. Table S2
    c. Table S3
    d. Table S4
    e. Table S5

### 1. Repeated Trust Game

In order to manipulate the type of social interaction experienced with the two confederates, a modified version of the Repeated Trust Game was used. In particular, the back-transfer behavior of the two trustees (confederates) was computer-controlled in the same way as in Rosenberger et al. 2020, so that participants always played with a cooperative and a non-cooperative trustee. Participants did not receive any priors about the cooperativeness of the trustees. In the first four trials, back-transfers of both trustees were fixed to 100% of the participant's investment in order to avoid ceiling or floor effects on the investment and to create a common prior about the trustees' back-transfers among participants. After the fourth trial, back-transfers of the cooperative trustee were, with an equal probability, either 100%, 150%, or 200% of the participant's investment. Every time the participant increased the investment towards the cooperative trustee in consecutive trials, the probability of 200% investment back-transfer increased by 10%, and the probabilities of the other two options decreased by 5% each. Back-transfers of the non-cooperative trustee were, with an equal probability, either 100%, 50%, or 75% of the invested amount. Every time the participant decreased the investment towards the non-cooperative trustee in consecutive trials, the probability of receiving the 50% investment back-transfer increased by 10%, and the probabilities of the other two options decreased by 5% each. Participants had to learn throughout the game to invest as much as possible in the cooperative trustee and as little as possible in the non-cooperative one, in order to maximize their profits at the end of the game. The two confederates played the cooperative or non-cooperative trustee randomly across participants. Prior to the task, participants received detailed written instructions in which they were also informed that their final profit in the game was paid at the end of the study. Regardless of the actual win, at the end of the session all participants received 5 € (in addition to the 25€ compensation for their participation in the study).

After the instructions and before the task, participants had to fill out control questions where they calculated profits for different scenarios. If the calculations were incorrect participants received additional verbal explanations until they fully understood the task. Participants (and confederates) also played five practice trials with the computer as a trustee and received additional in-game instructions to familiarize themselves with the layout of the task.

### 2. Validity of the cover story

In order to check if participants believed in the cover story, we examined their answers to the questions about credibility they filled out at the end of the study (e.g., “Did you notice something unusual during the space task?”, “What were your thoughts about the two other players?”). We therefore categorized participants in *believers* (participants who did not express any doubts about the experimental procedure) and *non-believers* (participants who expressed doubts about either the Repeated Trust game or the Interpersonal Space task procedures). We then investigated possible differences in the number of believers and non-believers across the two groups computing a chi-squared ($\chi^2$) test (Fisher’s Exact test). Lastly, we assessed the influence of non-believers on the behavioral data by computing the before mentioned ANOVAs using the “belief” variable as a covariate. The “belief” variable was a dichotomous variable, where 1 represented participants who believed in the task and 0 defined participants who expressed doubts about its veracity.

Overall, seven participants (five ASD) expressed doubts regarding the veracity of the experimental procedure. No significant relationship between group and veracity was detected ($\chi2$ (1) = 1.59, p = 0.41). When the belief information was added as a covariate to the main behavioral analyses, no changes in the results were observed. Moreover, the reported behavioral data analyses were also conducted after removal of these seven participants and showed no changes in the results.

### 3. Results from ANOVA performed at T1 (behavioral data)

To confirm that the results regarding interpersonal space *permeability* were not affected by using the data from the second run (T2; after the trust game), we performed the same mixed model ANOVA including the between-subjects factor Group (ASD, CTR) and the within-subject factors Step (1-5) and Trustee (cooperative, non-cooperative) on the data from the first run (T1) only. The analysis revealed a similar pattern of results as in the original analysis. In particular, we observed significant main effects of Group ($F_{1,38}$ = 6.19, $p$ = .017, $\eta^2_G$ = .045) and Step ($F_{1.69,64.1}$ = 72.04, $p$ < .001, $\eta^2_G$ = .55). No significant main or interaction effects of Trustee were found (all $F$ < .52, all $p$ > .51), confirming that, before the trust game, the comfort ratings were independent from the confederate approaching the participant.

### 4. Supplementary tables

*Table S1.* Means and SDs (in brackets) of Interpersonal Space task (comfort).

| | | ASD | | CTR | |
|---|---|---|---|---|---|
| **Interpersonal Space Task** | | Cooperative | Non-cooperative | Cooperative | Non-cooperative |
| Step 1 | T1 | 4.16 (4.19) | 4.02 (4.66) | 5.83 (3.56) | 5.50 (3.97) |
| | T2 | 3.88 (4.44) | 4.00 (4.15) | 5.17 (4.24) | 4.26 (4.27) |
| Step 2 | T1 | 3.46 (2.92) | 2.93 (3.32) | 4.79 (2.25) | 4.74 (2.48) |
| | T2 | 2.58 (2.86) | 1.30 (2.60) | 4.15 (2.40) | 2.98 (2.82) |
| Step 3 | T1 | 1.70 (3.69) | 1.47 (3.78) | 3.02 (3.23) | 3.07 (3.41) |
| | T2 | 1.53 (3.34) | -0.09 (3.89) | 3.44 (3.76) | 1.54 (4.15) |
| Step 4 | T1 | -1.40 (4.71) | -1.46 (5.09) | 0.54 (5.07) | 0.89 (4.92) |
| | T2 | 0.13 (5.06) | -1.55 (5.34) | 2.20 (5.41) | -0.11 (5.20) |
| Step 5 | T1 | -7.25 (3.49) | -7.66 (2.71) | -6.01 (3.99) | -6.11 (4.15) |
| | T2 | -6.66 (2.90) | -7.93 (2.72) | -5.13 (4.65) | -6.81 (4.10) |

*Table S2.* Means and SDs (in brackets) of the confederates' picture ratings (trustworthiness, fairness, intelligence, attractiveness) and Repeated Trust Game (investment).

| | ASD | | CTR | |
|---|---|---|---|---|
| **Picture Rating** | Cooperative | Non-cooperative | Cooperative | Non-cooperative |
| Trustworthiness T1 | 1.72 (3.34) | 1.78 (3.77) | 0.85 (4.31) | 2.77 (2.90) |
| Trustworthiness T2 | 4.28 (3.37) | -1.88 (4.81) | 4.56 (4.83) | -1.15 (5.62) |
| Fairness T1 | 1.95 (4.01) | 2.36 (4.32) | 2.56 (3.82) | 4.00 (3.14) |
| Fairness T2 | 5.41 (4.12) | -2.95 (4.84) | 6.36 (4.03) | -2.48 (6.07) |
| Intelligence T1 | 3.64 (3.01) | 2.50 (3.51) | 3.80 (2.60) | 3.94 (3.98) |
| Intelligence T2 | 3.78 (3.55) | 1.14 (5.10) | 4.96 (4.89) | 2.42 (5.55) |
| Attractiveness T1 | 0.45 (4.70) | 1.38 (4.41) | -0.73 (4.91) | 1.86 (5.50) |
| Attractiveness T2 | 0.64 (4.57) | -0.12 (5.47) | 0.15 (5.23) | 0.96 (5.63) |
| **Repeated Trust Game** | | | | |
| Investment | 8.01 (1.57) | 3.25 (1.52) | 7.95 (2.12) | 4.07 (1.66) |

*Table S3.* Coordinates used to create the mask used for small volume correction (SVC) in the main fMRI analyses.

| Region | Hemi | x | y | z | Reference |
|---|---|---|---|---|---|
| FFA | R | 34 | -65 | -16 | Foss-Feig et al., 2016 |
| | R | 36 | -53 | -19 | |
| | L | -38 | -65 | -18 | |
| | L | -38 | -51 | -18 | |
| hMT+/V5 | R | 40 | -67 | -3 | Foss-Feig et al., 2016 |
| | L | -42 | -79 | -3 | |
| Amygdala | R | 18 | 3 | 18 | Kennedy et al., 2009 |
| | L | -18 | -3 | 15 | |
| dIPS | R | 10 | -53 | 57 | Holt et al., 2014,2015 |
| | R | 25 | -51 | 52 | |
| | R | 35 | -41 | 57 | |
| | R | 30 | -33 | 44 | |
| | R | 10 | -57 | 59 | |
| | L | -19 | -70 | 40 | |
| | L | -17 | -57 | 59 | |
| | L | -34 | -44 | 56 | |
| | L | -12 | -52 | 56 | |
| | L | -22 | -47 | 57 | |
| | L | -23 | -51 | 61 | |
| | L | -14 | -49 | 59 | |
| PMv | R | 52 | -3 | 29 | Holt et al., 2014,2015 |
| | R | 55 | 5 | 29 | |
| | L | -51 | -3 | 27 | |
| | L | -55 | -2 | 35 | |
| | L | -50 | 2 | 5 | |
| | L | -52 | -4 | 43 | |

In order to create the mask for SVC in the main fMRI analyses, the listed coordinates were used to build 8-mm spheres, which were then combined in a single mask using the MarsBar SPM Toolbox. Hemi, hemisphere; x y z, MNI coordinates. FFA, Fusiform Face Areas; hMT+/V5, human middle temporal area; dIPS, dorsal Intraparietal Sulcus; PMv, ventral Premotor Cortex.

*Table S4.* Whole brain analysis results ($p < .05$ FWE corrected) in the computed contrasts of interest.

| Region | Hemi | Cluster K | Z | $p_{FWE}$ | x | y | z |
|---|---|---|---|---|---|---|---|
| CTR > ASD | | | | | | | |
| Not assigned | L | 874 | 6.0 | .000 | 0 | -92 | 4 |
| Not assigned | L | | 5.6 | .001 | -2 | -90 | 22 |
| Inferior Occipital Gyrus | R | 114 | 5.0 | .018 | 44 | -62 | 4 |
| Inferior Frontal Gyrus (p. opercularis) | R | 160 | 4.9 | .023 | 46 | 14 | 28 |
| Precentral Gyrus | L | 333 | 4.9 | .031 | -32 | -10 | 66 |
| Superior Frontal Gyrus | R | 338 | 4.8 | .034 | 6 | 12 | 56 |

| | | | | | | | |
|---|---|---|---|---|---|---|---|
| ASD > CTR | | | | | | | |
| Middle Occipital Gyrus | L | 322 | 4.9 | .021 | -18 | -100 | -2 |
| (CTR: (T2: FAIR > UNFAIR) > (T1: FAIR > UNFAIR)) > (ASD: (T2: FAIR > UNFAIR) > (T1: FAIR > UNFAIR)) | | | | | | | |
| No suprathreshold voxels | | | | | | | |
| (ASD: (T2: FAIR > UNFAIR) > (T1: FAIR > UNFAIR)) > (CTR: (T2: FAIR > UNFAIR) > (T1: FAIR > UNFAIR)) | | | | | | | |
| No suprathreshold voxels | | | | | | | |
| (CTR: FAIR > UNFAIR) > (ASD: FAIR > UNFAIR) | | | | | | | |
| No suprathreshold voxels | | | | | | | |
| (ASD: FAIR > UNFAIR) > (CTR: FAIR > UNFAIR) | | | | | | | |
| No suprathreshold voxels | | | | | | | |
| (T2: FAIR > UNFAIR) > (T1: FAIR > UNFAIR) | | | | | | | |
| No suprathreshold voxels | | | | | | | |
| (T2: UNFAIR > FAIR) > (T1: UNFAIR > FAIR) | | | | | | | |
| No suprathreshold voxels | | | | | | | |

Hemi, hemisphere; x y z, MNI coordinates; Z, z scores; Cluster K, number of voxels in associated cluster. Voxels were labelled based on MRIcron Jhulich template.

*Table S5.* BMA of PEB parameters: Intrinsic connectivity (*A* matrix) and direct input (*C* matrix).

| **A matrix** | | | | | |
|---|---|---|---|---|---|
| Source | Target | Commonalities | Group | Comfort | Group x Comfort |
| Amy | Amy | -0.65 | 0.34 | -0.05 | 0.00 |
| Amy | dIPS | 0.22 | -0.24 | 0.00 | 0.10 |
| Amy | FFA | 0.43 | -0.28 | 0.03 | 0.13 |
| dIPS | Amy | 0.00 | 0.13 | 0.00 | 0.05 |
| dIPS | dIPS | 0.14 | -0.11 | 0.00 | 0.00 |
| dIPS | FFA | -0.35 | 0.18 | -0.04 | 0.00 |
| FFA | Amy | 0.00 | -0.15 | 0.00 | 0.04 |
| FFA | dIPS | 0.19 | 0.00 | -0.04 | -0.03 |
| FFA | FFA | -0.29 | 0.00 | -0.07 | 0.00 |
| V1 | Amy | 0.07 | 0.00 | 0.00 | 0.00 |
| V1 | FFA | 0.17 | -0.02 | -0.01 | -0.02 |
| V1 | V1 | 0.47 | 0.00 | -0.04 | -0.02 |
| **C matrix** | | Commonalities | Group | Comfort | Group x Comfort |
| | | 0.65 | 0.00 | 0.00 | 0.00 |
